# Polee: RNA-Seq analysis using approximate likelihood

**DOI:** 10.1101/2020.09.09.290411

**Authors:** Daniel C. Jones, Walter L. Ruzzo

## Abstract

The analysis of mRNA transcript abundance with RNA-Seq is a central tool in molecular biology research, but often analyses fail to account for the uncertainty in these estimates, which can be significant, especially when trying to disentangle isoforms or duplicated genes. Preserving uncertainty necessitates a full probabilistic model of the all the sequencing reads, which quickly becomes intractable, as experiments can consist of billions of reads. To overcome these limitations, we propose a new method of approximating the likelihood function of a sparse mixture model, using a technique we call the Pólya tree transformation. We demonstrate that substituting this approximation for the real thing achieves most of the benefits with a fraction of the computational costs, leading to more accurate detection of differential transcript expression.

**Availability:** The method is implemented in a Julia package available from https://github.com/dcjones/polee

## 1 Introduction

The past decade has seen RNA-Seq become a central tool in molecular biology research. Along the way there have been numerous methods developed to analyze this data. We will propose an entirely new methodology based on likelihood approximation, which enables inference on full probabilistic models that would otherwise quickly grow intractable. In preparation, we will first give a brief overview of notable approaches to RNA-Seq transcript and gene quantification, to give some sense of where this new method fits in.

Gene/transcript quantification is not the only application of RNA-seq. Most prominently, the technology has been used to discover and annotate new transcripts, either by de novo assembly [Robertson et al., 2010, Grabherr et al., 2011, Haas et al., 2013], or processing reads aligned to a reference genome sequence [Trapnell et al., 2010, Guttman et al., 2010, Pertea et al., 2015]. Other applications include detection of fusion transcripts [Kumar et al., 2016] and RNA editing [Peng et al., 2012]. We will largely ignore these uses of RNA-seq to focus squarely on quantification. For the most part, we will assume that a suitable reference genome sequence and transcript annotations are available (or simply transcript sequences).

Because of alternative splicing, alternative transcription start and termination sites, and paralogous genes, transcripts often have a degree of sequence similarity that renders short reads ambiguous. Short read RNA sequencing in transcriptionally complex organisms thus produces a mixed signal. A broad distinction that must be drawn among quantification methods is between those that avoid trying to deconvolute these mixed signals and those that embrace deconvolution. A fundamental assumption of most RNA-seq analyses is that transcript expression is proportional in expectation to the number of reads observed from that transcript (when sample specific effects are accounted for). As Mortazavi et al. [2008] notes, read counts were observed to be “linear across a dynamic range of five orders of magnitude in RNA concentration.” Yet reads of ambiguous origin cannot be trivially assigned to transcripts. Estimating transcript expression thus necessitates either ignoring ambiguous reads, explicitly assigning them to transcripts, or otherwise implicitly considering the space of possible assignments.

Transcript expression is not the only possible quantity of interest, and many methods have found success in posing alternative inference problems that avoid dealing with read ambiguity and deconvolution altogether. These methods are often simpler and more efficient than deconvolution methods, so with good reason they remain popular, yet they are fundamentally limited in scope. DEXSeq [Anders et al., 2012] approaches the problem by considering exon usage. JunctionSeq [Hartley and Mullikin, 2016] and LeafCutter [Li et al., 2018] are both methods that specifically focus on reads crossing splice junctions, with the goal of detecting changes in usage.

Some count-based approaches to gene expression assume that genes (however defined) are reasonably unambiguous, so that the ambiguous reads that are present are few and relatively inconsequential. However, many methods do deal directly with the deconvolution problem, which will be our interest. Though there are non-probabilistic approaches to this problem, including methods using network flow for transcript quantification [Montgomery et al., 2010] and assembly [Kannan et al., 2016], and integer programming Lin et al. [2012], the probabilistic approach has been by far the most common and is the one we will focus on here.

The common probabilistic approach to the transcript quantification problem is to treat transcripts as inducing distinct probability distributions over reads (or read pairs). The experiment as a whole can then be thought of as a mixture model, in which the goal is to infer relative transcript expression (i.e., mixture coefficients). More explicitly, given a set r of m reads, and n annotated transcripts, we define a probability function p_j_ over possible reads, for every transcript j. The likelihood for a relative expression vector x ∈ Δ^n−1^ (here Δ^n−1^ is the open unit (n − 1)- simplex, i.e., the set of all vectors of length n, with positive entries summing to 1) is

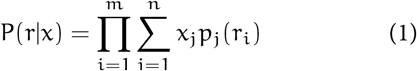

There are many issues surrounding the question of how best to model each transcript read distribution p_j_. RNA-Seq protocols involve many steps like fragmentation, reverse transcription, amplification, and fragment size-selection which each influence the observed distribution of reads. Assuming reads to be uniformly distributed across a transcript, subject to some fragment length distribution, is the most straightforward model, but some success has been had in building more accurate models that capture positional and sequence-specific biases (see for example [Hansen et al., 2010, Li et al., 2010, Roberts et al., 2011, Jones et al., 2012]). We will set these issues aside for now and assume that we have some agreed upon model.

Relative transcript expression for an individual RNA-Seq sample is not typically interesting on its own. RNA-Seq experiments are nearly always concerned with detecting transcriptional changes between groups of samples. From a Bayesian perspective, we would like to build a model of transcriptional changes among k samples consisting of sets of reads r^(1)^, …, r^(k)^, with model parameters θ (e.g. effect sizes, pooled means, or latent space encodings), then consider the posterior distribution P(θ|r^(1)^, …, r^(k)^) ∝ P(r^(1)^, …, r^(k)^|θ)P(θ).

If we adopt the likelihood function in Equation 1, then r^(i)^ is independent of all other variables when conditioned on x^(i)^. In typical models expression vectors x^(i)^ will also be mutually independent when conditioned on the model parameters θ, so we can write this posterior distribution in terms of the latent expression vectors x^(1)^, …, x^(k)^, treating them as nuisance parameters.

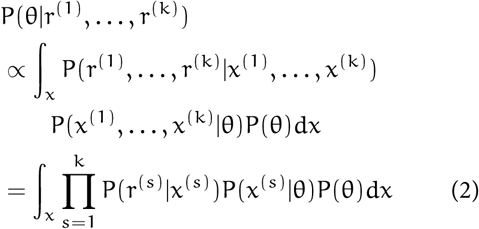

This is all very straightforward but presents some practical problems. To evaluate the likelihood functions, some information about each unique read must be stored (typically in a sparse matrix where entry i, j corresponds to the probability assigned to the ith read by the jth transcript distribution). This translates to hundreds of megabytes to several gigabytes per sample. Estimating the posterior for moderately large experiments requires either a great deal of memory or cycling read data in and out of memory, as is done in stochastic gradient methods.

In practice, this kind of textbook model is often short-circuited. Point estimates are made for transcript expression vectors x, usually by maximum likelihood

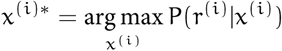

Then these are plugged into the full model, forming an alternative posterior distribution

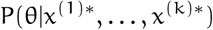

This model adopts the (false) assumption that these expression values are observed, substituting them for what is actually observed: the reads. Non-Bayesian models have the same issues, and often resort to the same two-step approach. When estimates are treated as observations, the uncertainty of these estimates is flushed from the analysis, artificially inflating the certainty of the end result. This two-step approach can be thought of as an approximation of the desired model, but one that captures none of the uncertainty of read assignments.

A better two-step approach has been implemented by Pimentel et al. [2017] in their work on *sleuth*. Sleuth uses bootstrap samples to estimate uncertainty in maximum likelihood estimates. Incorporating these variance estimates into regression models, they show substantial improvements in accuracy when calling differential expression, particularly at the transcript level. Bootstrap methods do have limitations, though. Estimates of variance are guaranteed to converge asymptotically to the true value with enough reads, but this leaves lightly sequenced loci with potentially unreliable estimates.

Are there better approximations? A similar approach to sleuth could be taken but with MCMC samples. Unlike the bootstrap, MCMC estimates of variance would not be contingent on sufficiently deep sequencing of a locus, so may be more reliable when capturing uncertainty of low expression isoforms. But samples have limited usefulness. For example, powerful probabilistic programming languages have become an increasingly popular way to implement models, but efficient inference usually relies on variational inference or some form of Hamiltonian Monte Carlo. To make use of these tools, we would want a compact approximation of the likelihood function that can be efficiently evaluated and differentiated. In the next section we propose such a solution.

## 2 Approximate likelihood

In this section we develop a novel approximation for the RNA-Seq likelihood function. Because the likelihood function for a full experiment factors into per-sample likelihood functions (as in Equation 2), this approximation can be built one sample at a time. Once fit, evaluating and sampling from the approximation is orders of magnitude faster than using the likelihood function. Substituting this approximation for the real thing can make inference on full probabilistic models, with billions of reads, tractable on even modest computers.

There has not been much work exploring the idea of approximating the RNA-Seq likelihood function, but one notable exception is Zakeri et al. [2017], who developed an approach in which reads that have similar values assigned by p_j_(·) for every transcript j are treated as equivalent and combined in Equation 1. This can significantly improve efficiency of the likelihood function with only a moderate decrease in fidelity. What we propose goes further, reducing the likelihood function into an exceedingly efficient constant time and space function, with only slight reductions in accuracy.

### 2.1 Approximating likelihood with variational inference

Variational inference is usually presented as a means of estimating an otherwise intractable posterior distribution. Given a distribution function p, and a family of distributions q(·; ϕ) parameterized by ϕ, we fit q to p, given some data y, by choosing ϕ to minimize the Kullback-Leibler (KL) divergence,

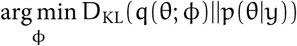

A key feature of this method of inference is that q does not depend directly on y, so that once ϕ is optimized, the approximate probability can be evaluated without the data, only retaining the typically much smaller parameter vector ϕ.

This suggests a solution to the issue of building large joint models. Instead of using variational inference to approximate a posterior distribution, we can use it to separately approximate the likelihood of each sample. By substituting an approximation q(x; ϕ) for each sample’s likelihood function P(r|x), we can capture the likelihood with some fidelity without having to keep the RNA-Seq reads in memory. This would allow us to build a very large model, encompassing hundreds or thousands of samples, that can be run on laptops or meager servers.

Of course, the likelihood is not a distribution over expression vectors x but over reads r, so the KL divergence is not well-defined here. But in models making use of the likelihood function, multiplicative constants are generally irrelevant, so our approximation only has to be proportional to the likelihood. To bring the machinery of variational inference to bear, instead of approximating likelihood directly, we approximate a normalized likelihood function, mathematically equivalent to a posterior distribution under a uniform prior. We will denote with 𝒫(x|r) this normalized likelihood function, defined simply as

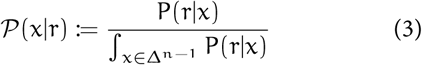

In Figure 1, some illustrative examples of the approach we develop in the following sections are shown. We can see that capturing the dependence structure of 𝒫 requires careful selection of the distribution family being used, but it often possible to do so very accurately. In simple cases lacking any read ambiguity, the likelihood is proportional to a Dirichlet distribution, but this model is inadequate for cases where reads are compatible with multiple transcripts. The model we propose can perfectly capture the cases of zero read ambiguity (see Appendix B.2), but is strictly more expressive, and able to capture more complex dependence structures while being similarly efficient (i.e. linear time and space in the number of transcripts).

**Figure 1:**
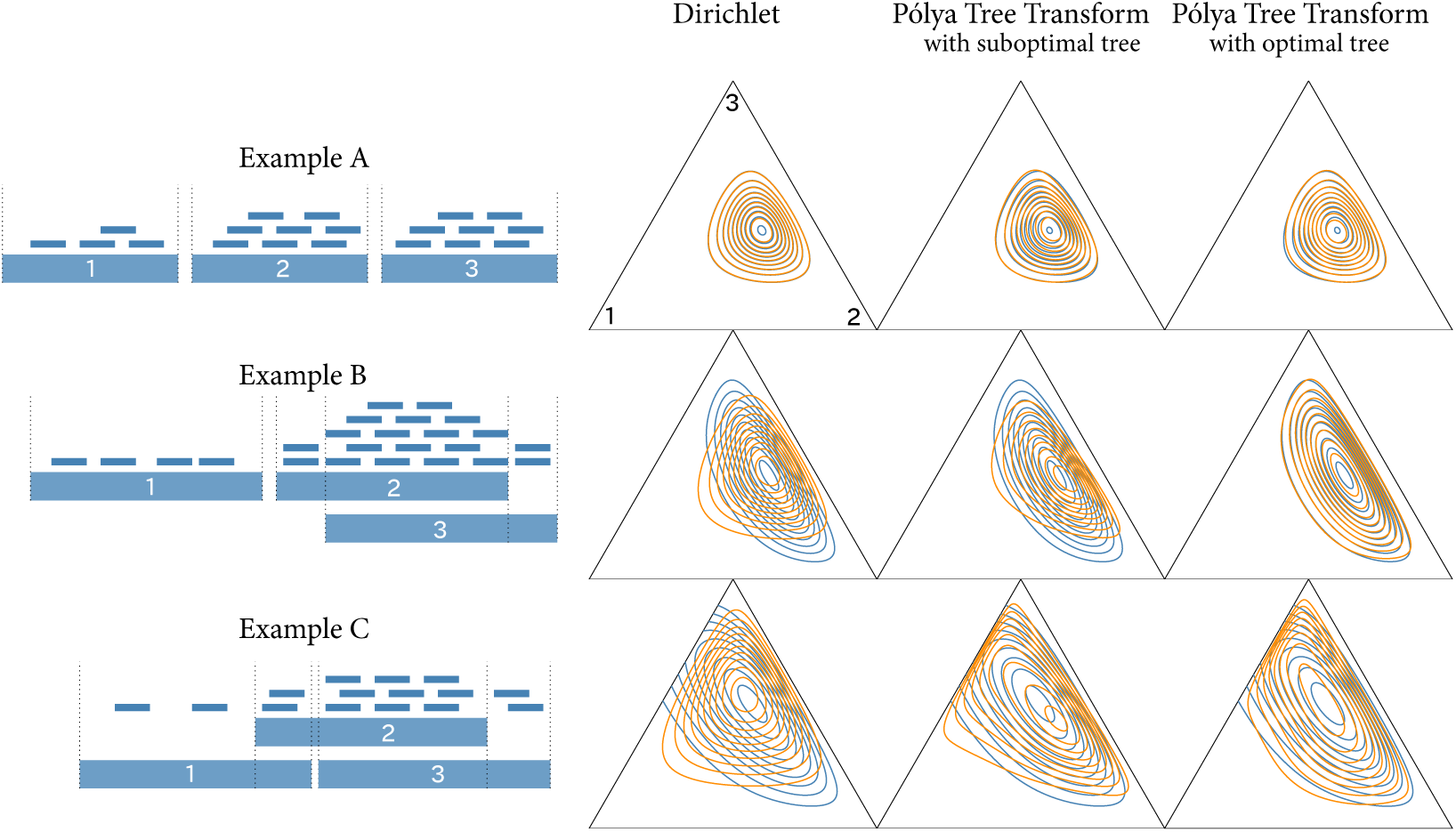
Three contrived RNA-Seq examples are shown on the left. On the right, in blue, contours of the their likelihood functions are plotted, and in orange, various approximations made by minimizing KL divergence. The approximation we propose, based on what we call the Pólya tree transformation, is shown in the second and third columns. This transformation is defined in terms of a tree. Choosing the right tree plays a large role in how well the approximation fits. In example A, every read can be unambiguously attributed to a single transcript. In this easy case, a Dirichlet distribution is perfectly proportional, and the approximation, regardless of the tree, is a near perfect fit. In examples B and C, reads cannot by unambiguously assigned. Here, the proposed approximation remains a good fit, provided the right tree is selected. Using the approximation in place of the exact likelihood function can enable dramatically more efficient inference.

#### 2.1.1 Practical benefits to likelihood approximation

Our approach to approximation is unusual: factoring the likelihood, then fitting a proportional approximation to each factor. Though it could be used in other settings, it is particularly well suited for RNA-Seq, compared to other possible approaches.

The obvious alternative for tractable inference is simply to use variational inference on the posterior we are actually interested in. That is, if we have a model of, say, differential expression, with parameters θ, we want to approximate the intractable posterior P(θ|r). For a large experiment, all of the reads r will not fit in memory, but this problem is amenable to stochastic variational inference, or SVI [Hoffman et al., 2013]. In SVI, batches of data are subsampled to update estimates of “local” latent parameters (transcript expression estimates x^(i)^ for each sample i), before updating “global” latent parameters (θ).

Because SVI algorithms must cycle many gigabytes of sequencing data in and out of memory to repeatedly compute stochastic gradients, they are likely much less efficient than likelihood approximation. Two additional considerations further increase the latter’s desirability.

First is reusability. Large RNA-Seq experiments can be rich with insight, and lend themselves to multiple analyses. Differential expression, differential splicing, clustering, and dimensionality reduction are separate tasks that might be carried out on the same data, each with its own model, each representing a separate inference task. For these tasks, likelihood approximations must be made only once, after which they can be reused over an over. Amortized over every iteration of every analysis typically run on a dataset, likelihood approximation is far more efficient than other approaches to tractable inference.

Second, likelihood approximation can obviate some of the cumbersome data transfer and storage issues with RNA-Seq. High throughput sequencing produces huge datasets, which must be stored and transferred to collaborators, which can mean waiting on long downloads or exchanging hard drives. Our approach to approximated likelihood, on the other hand, summarizes all expression information for a sample using only a few megabytes per sample. The 1461 brain samples produced by the GTEx project [GTEx Consortium, 2013], are reduced to 7.8GB of likelihood approximation data. The most compact exact representation of the likelihood function for this experiment would be about 1.5TB, often a prohibitively large amount of data to keep in memory.

### 2.2 Designing an approximation

Because RNA-Seq data is compositional, our approximating family of distributions q(x; ϕ) must be defined over a simplex Δ^n−1^. Without going into the mathematical details (see Appendix B.3), the common approach to optimizing the KL-divergence additionally requires a distribution that can be expressed as a deterministic bijection of a random variable drawn from some fixed distribution (a technique termed the “reparameterization trick” [Salimans and Knowles, 2013, Kingma and Ba, 2014]).

The Dirichlet distribution tends to be the default simplical distribution, but besides being insufficiently expressive for our goals (see Figure 1), it is not efficiently computable in terms of reparameterization. Instead, we consider transformations of other distribution families onto the simplex. For this to work we need a bijection T : ℝ^n−1^ → Δ^n−1^ (or perhaps T : (0, 1)^n−1^ → Δ^n−1^), with an efficiency computable Jacobian determinant.

#### 2.2.1 Compositional data analysis transformations

The compositional data analysis literature has traditionally been concerned with how best to transform data to

and from the simplex. A number of Δ^n−1^ → ℝn−1 transformations have been proposed, some with inverses. When used to transform a normal distribution, for example, these can induce useful simplex distribution families. Three such bijective transformations are explored here as possible candidates to define a suitable distribution.

The first common approach is the additive log-ratio transformation.

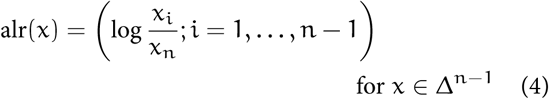

The alr is typically defined, as it is here, with the divisor x_n_, but the vector can be permuted to make any element the divisor. In some settings it makes sense to choose a particular element as the reference for interpretability; for example we might choose x_n_ to be the expression of a housekeeping gene. When searching for the best fitting approximation, there is not an obvious choice. If y ∼ Normal(µ, Σ), then the distribution induced by alr^−1^(y) is sometimes referred to as a *multivariate logit-normal distribution*.

The second useful transformation defined by Aitchison is the *multiplicative log-ratio transformation*

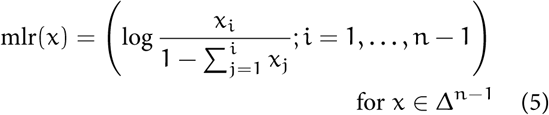

This transformation can be best understood using a sequential stick-breaking metaphor. If a stick is broken into n pieces, and x_1_, …, x_n_ give the size of each piece, in proportion to the whole, then mlr(x) gives the log-ratio between each piece and the remaining length of the stick, if the stick were broken one piece at a time. We will return to this stick-breaking metaphor shortly.

The probabilistic programming language Stan [Carpenter et al., 2016] implements essentially this transformation as a general purpose variational approximation to distributions on the simplex as part of its Automatic Differentiation Variational Inference approach [Kucukelbir et al., 2017].

The third and most modern approach from compositional data analysis is the *isometric log-ratio* transformation [Egozcue et al., 2003]. The mathematical details of this are too involved to go into here, but it has a number of desirable features. Most significantly it is, as the name implies, an isometry, or distance preserving transformation. So the Euclidean distance between two transformed points ∥ilr(x)−ilr(y) ∥ = d_a_(x, y), where d_a_ is the *Aitchison distance*, whose definition will a lso be omitted here. That geometry is preserved by ilr ends up making certain statistical procedures interpretable on the simplex where they might otherwise not be.

Figure 2 gives some intuition for how these transformations operate.

**Figure 2:**
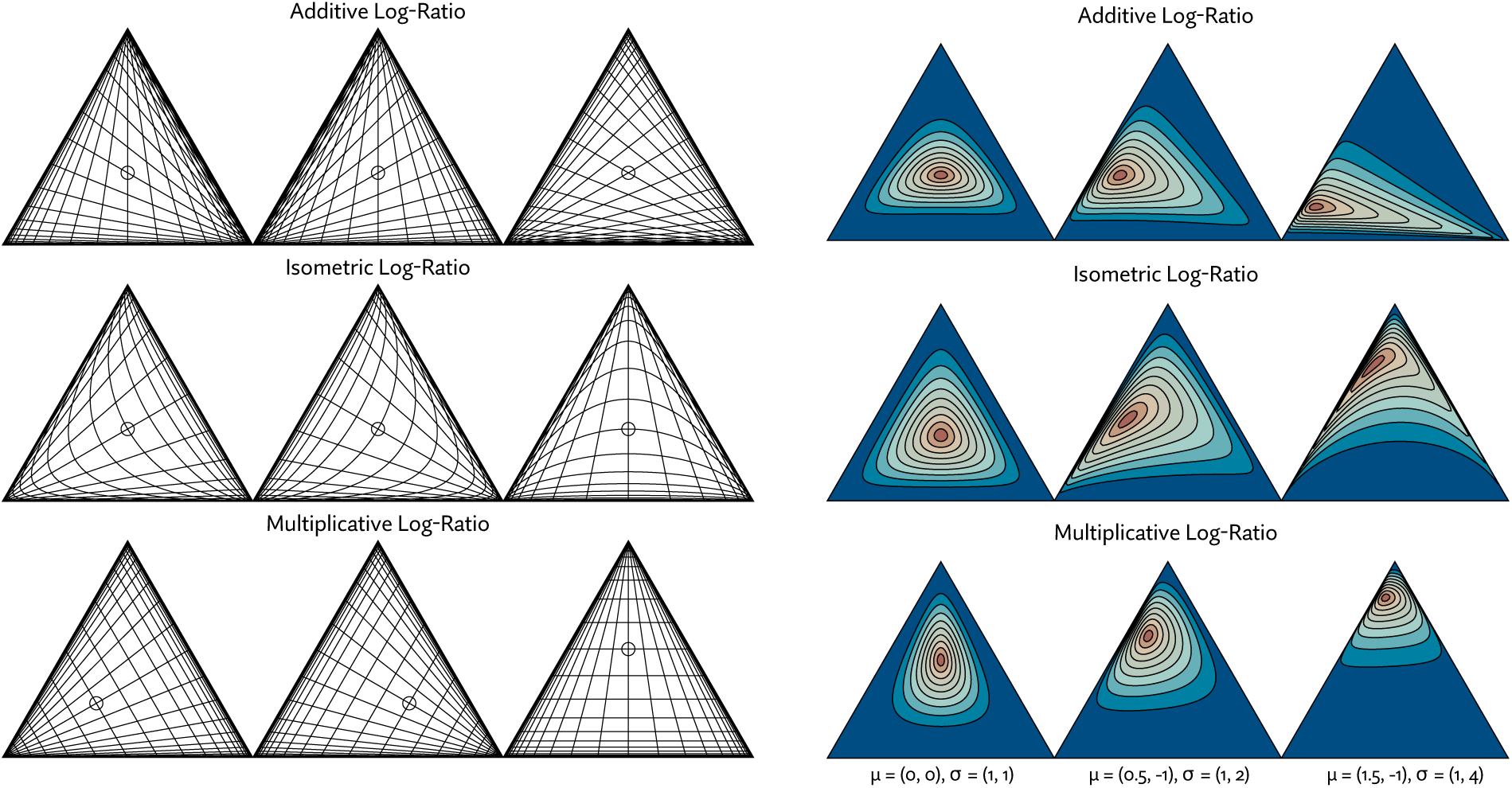
On the left, Cartesian grid lines are transformed onto Δ^2^ using three classes of transformations: additive log-ratio, isometric log-ratio, and multiplicative log-ratio. Grid lines are spaced evenly at a distance of 0.5 in Euclidean space and the point (0, 0) is marked with a circle. Each transformation has variations shown in the columns. The variations are formed by choosing a different denominator, basis, or permutation for alr, ilr, and stick breaking, respectively. On the right, various parameterizations of a 2-dimensional multivariate normal distribution, with a diagonal covariance matrix, are mapped onto the simplex with alr, ilr, and mlr (for each, the furthest right of the three variants shown on the left). The choice of transformation has a dramatic effect on the resulting distribution.

#### 2.2.2 The Pólya tree transformation

To revisit the stick-breaking metaphor, it is often easiest to consider generating a vector x ∈ Δ^n−1^ by starting with a stick of length 1 and breaking it n − 1 times in sequence.

Let y_i_ ∈ (0, 1) represent the proportion of the remaining stick to break off on the ith break. Then x ∈ Δ^n−1^ can be produced from y ∈ (0, 1)^n−1^ with

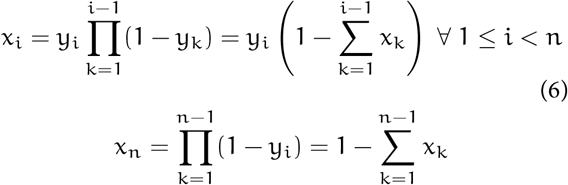

The equivalence between the product and sum form shown here may not be immediately obvious, but is easy to show by induction [Halmos, 1944], or by considering that 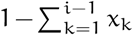 is the length of the stick remaining after_k_ *i* − 1 breaks.

Stick-breaking metaphors like the one used to describe the mlr transformation have a long history. In an early example, Halmos [1944] explores a stick breaking distribution in which each y_i_ is uniformly distributed in (0, 1) using the motivation of distributing a pile of gold dust to n beggars in sequence. The possibility of y_i_ being drawn independently from arbitrary distributions is also briefly considered. In recent literature, stick breaking occurs most commonly in descriptions of the Dirichlet process, which can be formulated as an infinite stick breaking procedure in which the breaks y_i_ are Beta distributed variables [Sethuraman, 1994]. The resulting stick sizes are then used to weight draws from a base distribution. When n → ∞ in Equation 6, and each y_*i*_ is i.i.d. Beta(1, θ) distributed, for some, the resultin stribution family is commonly denoted GEM(θ), after Griffiths, Engen, and McCloskey (see Pitman [2002]). This notion of a stick breaking prior was generalized by Ishwaran and James [2001] to, among other things, include finite stick breaking distributions.

Khan et al. [2012] brings up an issue often ignored in these sequential stick-breaking procedures. They point out that the model represents a kind of decision boundary between every category i and the n − i categories that follow it in the process, so a particular ordering may fit the data poorly if no such boundary naturally exists. Zhang and Zhou [2017] take up this issue in a more serious way, demonstrating classification problems with as few as three categories that show dramatic differences in performance depending the permutation of those categories in the stick breaking process. They go on to propose models in which category permutations are inferred along with regression coefficients when performing multinomial logistic regression.

The second key insight that is sometimes neglected is that there are other ways of breaking a stick. Rather than breaking pieces off the stick and setting them aside, we might keep and recursively break both of the resulting pieces. This can be thought of as *hierarchical stick breaking*, as opposed to common *sequential stick breaking*. To define a transformation onto Δ^n−1^, we must always end up with n pieces, so n − 1 total breaks must still be made. Under these restrictions, breaks in a hierarchical stick-breaking scheme must occur according to a full binary tree with n leaves (i.e., where every node is either a leaf or has two children).

With these two insights, we have a space of possible stick-breaking transformations along the lines of Aitchison’s mlr, but considering not just the n! permutations of the stick breaking process, but also the 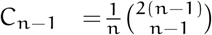 (the (n−1)st Catalan number) possible full binary trees with n leaves, resulting in a family of 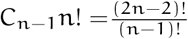 possible transformations.

This notion of hierarchical stick-breaking bears some resemblance to the hierarchical softmax transformation [Goodman, 2001], a technique used in some natural language models, but differs critically in that hierarchical soft-max is not bijective and is used purely as a means of accelerating inference. More closely related are Pólya tree distributions

[Lavine, 1992, 1994, Mauldin et al., 1992], which are also defined in terms of a (not necessarily finite) binary tree, in which each split or break is drawn from a Beta distribution. Due to this similarity, and apparently lacking any existing terminology, we refer to this family of hierarchical stick-breaking transformations as *Pólya tree transformations*. The software we implement to apply this method we call *Polee* (a portmanteau of “Pólya tree”).

#### 2.2.3 Tree topology heuristics

Though Zhang and Zhou [2017] were able to effectively optimize over stick-breaking topologies, at most 11 labels were used, and only permutations were considered. We would have a much harder time sampling over the possible configurations of a Pólya tree transformation. To avoid an exhaustive exploration of the space of trees we must either find an adequate heuristic with which to choose a tree, prior to optimizing parameters, or pursue optimization metaheuristics (e.g., simulated annealing or genetic programming) that could explore a subset of the space in a guided way. The latter may be possible, but each topology is accompanied with an entirely new set of parameters that must be optimized. Optimizing both concurrently is unlikely to be practical.

Fortunately, there are reasonable approaches we may take to choosing a topology ahead of optimization. RNA-Seq is typically a sparse mixture model. Most reads are compatible with only a small number of annotated transcripts. Because of this, the problem displays subcompositional independence: knowing the mixture of isoforms expressed in one gene tells us nothing about the mixture of isoforms expressed in another, if the genes share no reads. This suggests that the transformation might be oriented in a way to try to capture this structure.

Aitchison [1986] discusses the concept of subcompositional independence, as well as several other notions of independence on the simplex. There, tests for independence are proposed, but they necessitate estimating the full covariance matrix, an intractable task for a large n. Instead we pursue the idea of capturing a similar subcompositional independence structure by using hierarchical clustering as a heuristic. Each transcript is represented by its set of compatible reads. We then cluster greedily choosing the maximum Jaccard index (i.e. the size of the intersection divided by the size of the union) at each step. Transcripts, or sets of transcripts, that share a large proportion of their compatible reads have a have a higher Jaccard index, and thus their common ancestor is placed lower in the tree. In an ideal scenario, this constructs a tree that encodes a distribution family that has a similar independence structure to that of the real likelihood function.

In cases of complete subcompositional independence, where no reads are shared between transcripts, the likelihood function in Equation 1 is proportional to a Dirichlet distribution. In Appendix B.2 we show that any Pólya tree transformation can exactly fit any Dirichlet distribution of the same dimensionality, if applied to appropriately chosen Beta distributed random variables.

#### 2.2.4 Choosing a base distribution

Given a transformation onto the simplex, we now need to choose the distribution that will be transformed. The Pólya tree transform has been described described here as a (0, 1)^n−1^ → Δ^n−1^ transformation. Most of the compositional data analysis transforms take the form ℝ^n−1^ → Δ^n−1^. We can always use the logit function (or its inverse) to move between the two so this distinction is insignificant. We would like then to find a distribution over ℝ^n−1^ or (0, 1)^n−1^. To minimize the parameter space, each element will be considered independent (a common approach refereed to as “mean field variational inference”).

We are limited to distributions that lend themselves to the reparameterization trick. The two distributions we will consider are the normal (or logit-normal) distribution and the Kumaraswamy distribution [Jones, 2009], which is qualitatively similar to the Beta distribution but, unlike the Beta distribution, can be easily expressed as a transform of a uniform distribution.

Lastly we explore a further transformation of the normal distribution using a parameterized sinh-arcsinh transformation of the following form

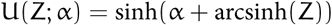

This technique, explored by Jones and Pewsey [2009] along with a two-parameter version, provides an analytically convenient way to add a parameter controlling skewness to a distribution. Where Z ∼ Normal(0, 1), we use

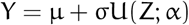

as our fully reparameterized distribution. We will refer to the resulting distribution as a *skew-normal* distribution, though other distribution families also go by this name [Azzalini, 1985, Hosking and Wallis, 2005].

## 3 Results

To demonstrate the usefulness of this approach, we undertook four analyses. In the first two subsections we show that the approximation is a good fit to the likelihood function of actual RNA-Seq datasets, and that it can be efficiently fit and evaluated. Then we focus on demonstrating that it can improve the detection of differentially expressed transcripts, using an existing simulation benchmark, and then separately using real data from the GTEx project.

### 3.1 The Pólya tree transform improves the goodness of fit to likelihood marginals

To assess the fit of our likelihood approximation, in comparison to various alternatives, we evaluated the fit of transcript marginal densities using Wilcoxon signed-rank tests. For every transcript in a set of annotations, 1000 samples were drawn from a Gibbs sampler (representing the exact likelihood function) and the same number of samples were drawn from the approximated likelihood. The signed-rank test was run, for each transcript, producing a p-value.

The Gibbs sampler used 8 randomly initialized chains. Each was burned-in for 2000 iterations, then 25,000 samples were generated and every 25th was saved for analysis, producing a total of 1000 samples. We found these settings sufficient for marginal distributions to have converged for a vast majority of transcripts, where convergence was measured by comparing within-chain and between-chain variance according to the procedure described by Gelman and Rubin [1992].

Sampling from the approximated likelihood is much simpler: random vectors are drawn from a Normal(0, I) distribution, then transformed using each of the transformations discussed, specifically the sinh-asinh transformation, shifting and scaling, the logistic transformation, and finally the Pólya tree transformation.

If an approximation is a perfect fit, we would expect to see a uniform distribution of p-values. Imperfect approximations will yield p-value distributions that are increasingly skewed towards smaller numbers. The greater the tendency towards small p-values, the worse the overall fit. Figure 3 shows the results of this test using a mouse brain sample taken from Li et al. [2017] (accession number PR-JNA375882), and 138,930 transcripts from the Ensembl 95 annotations [Ensembl, 2018]. To provide some intuition of the correspondence between p-value and fit, a number of examples with low p-values are plotted in Figure 4.

**Figure 3:**
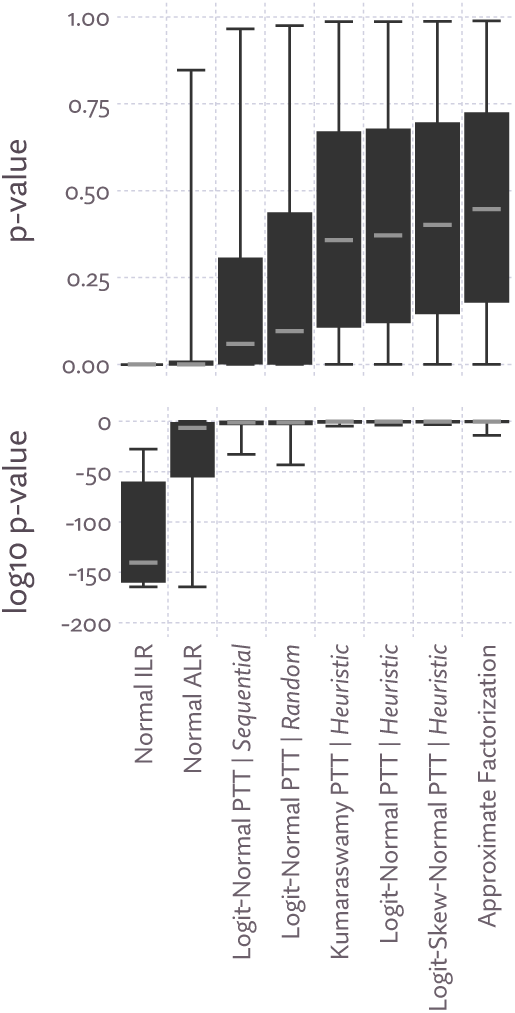
Distributions of p-values evaluating the agreement between 1000 samples from the Gibbs sampler and 1000 samples drawn directly from various approximations of the likelihood function for a mouse brain sample from Li et al. [2017]. P-values were computed using the Wilcoxon signed-rank test. Here, “PTT” is the Pólya tree transform, which was tried with several distribution families and tree building rules (“sequential”, “random”, and hierarchical clustering, labeled “heuristic”). The ilr and alr transformations are defined in Section 2.2.1, and “approximate factorization” is the approximation scheme proposed by [Zakeri et al., 2017]. Boxplots are drawn with upper and lower whiskers corresponding to the 99th and 1st percentile, respectively.

**Figure 4:**
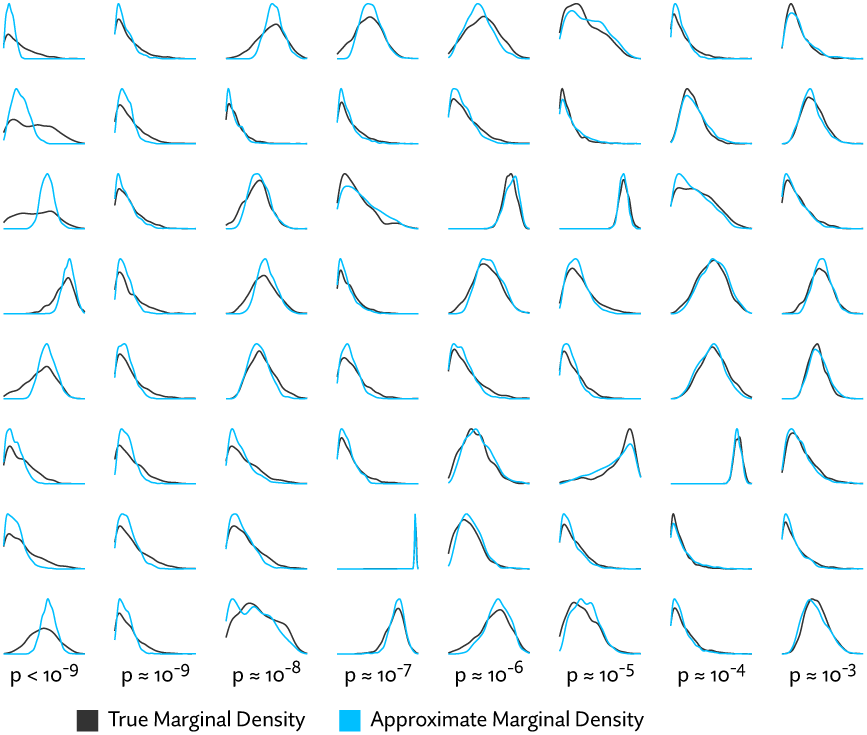
Kernel density plots of exact and approximate marginal densities for examples with small p-values. Columns correspond to p-values of decreasing powers of 10, while rows are randomly selected examples with approximately that p-value, giving a sense of how p-value corresponds to goodness of fit. It can be seen that a very small p-value does not necessarily correspond to a catastrophic failure of the approximation. The plots shown were generated from a mouse liver sample from Li et al. [2017] with approximate densities using logit-skew-normal Pólya tree transform distribution, using the hierarchical clustering heuristic to choose the tree topology.

We see that the approximating distribution family matters a great deal. Distributions based on the Pólya tree transformation offer a dramatically better fit than the traditional compositional data analysis transforms alr and ilr, and we see that the heuristic tree construction is a sharp improvement over a random tree topology. The common sequential stick-breaking approach (equivalent to the mlr transformation) is seen to be the worst approach to stick-breaking, perhaps in part because such a long chain of dependent breaks is numerically less stable and thus more difficult to fit to the target likelihood. As no attempt was made to optimize over permutations, it is also possible sequential stick breaking could be improved with the right permutation heuristic.

On this test, the approximate factorization approach proposed by Zakeri et al. [2017] also performs very well, offering a slight median improvement over the Pólya tree transformation. In Figure 5 we see that this comes at some cost. Though far more efficient in time and space that the un-approximated likelihood function, it does not match the low constant time and space of the Pólya tree transformation. Exact factorization (labeled “Factorization” in the figure), has a negligible effect, as few read pairs, even with 100 million reads, are completely identical.

**Figure 5:**
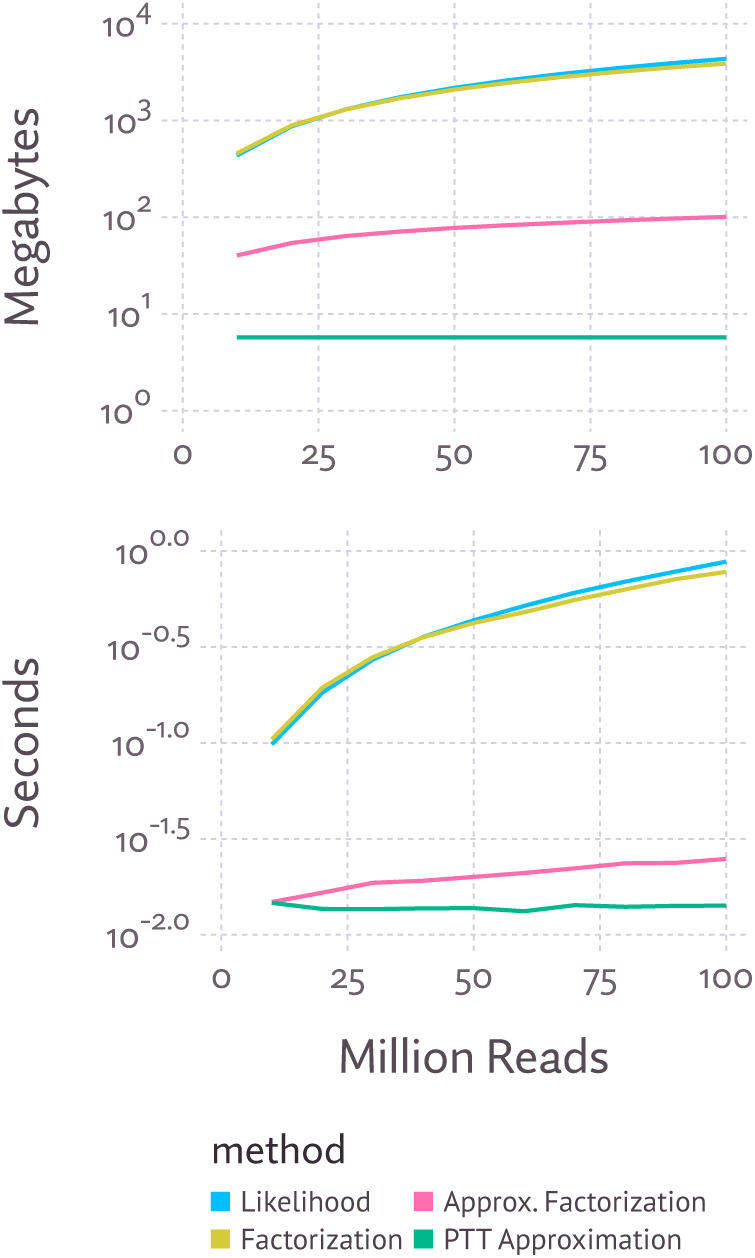
A comparison of the space and time needed to evaluate the likelihood function and its approximations. A GTEx sample (SRR1340508) was subsampled to, and its likelihood evaluated for, increasing numbers of reads. The upper plot shows the memory in megabytes necessary to evaluate the function, the lower plot, the time in seconds necessary to evaluate the function once.

This results holds across the other samples from Li et al. [2017] (Appendix A), but to ensure that the approximation is broadly effective and not somehow tuned to these particular datasets, we also evaluated logit-skew-normal Pólya tree transform distribution with heuristic tree topologies on a variety of of other RNA-Seq datasets, spanning a number of species with varying transcriptional complexity. We see that likelihood approximation has remarkably consistent performance across these samples (Table 1). Curiously though, the approximation appears to fit moderately better with a larger number of transcripts. The median p-value for both the human and mouse samples is approximately 0.4, and for the yeast sample, 0.28. Likely this is due to a smaller proportion of the transcripts being expressed in species with extensive alternative splicing.

**Table 1:** Auxiliary datasets used to evaluate the consistency of likelihood approximation performance, as was done in Figure 3. Median p-values are all for the proposed approximation (labeled “Logit-Skew-Normal PTT|Heuristic” in Figure 3). Numbers of transcripts listed are those annotated in version 90 of Ensembl. Accession numbers for these samples, from top to bottom, are: SRR453566, SRR030231, SRR065719, SRR023546, and SRR896663.

| Species | Num. Transcripts | Med. p-value |
| --- | --- | --- |
| S. Cerevisiae | 7126 | 0.28 |
| D. Melanogaster | 34749 | 0.37 |
| C. Elegans | 58941 | 0.35 |
| M. Musculus | 131195 | 0.40 |
| H. Sapiens | 200310 | 0.40 |

### 3.2 Estimating and sampling from approximated likelihood can be faster than bootstrap sampling

The procedure for sampling expression vectors from an approximated likelihood function is to simply generate a random vector from a Normal(0, I) distribution, and apply the transform, which is O(n) where n is the number of annotated transcripts. Because samples are so cheap to generate, for a sufficiently large numbers of samples, it outperforms not only MCMC approaches, but also the extremely fast bootstrap approach implemented in kallisto [Bray et al., 2016].

To locate the break even point, we recorded the overall time needed to generate increasingly large numbers of samples with both methods. In both approaches these times include the necessary initialization time. Kallisto uses its own pseudoalignment algorithm, while polee takes as input existing alignments. To compare on equal ground, we exported alignments generated by kallisto and used them as input into polee. Added to the polee timings is the time kallisto took to generate these alignments, the time it took to approximate the likelihood function, and to generated the requested number of samples. Both methods were run on 8 cpu cores.

From the results shown in Figure 6, we see that past about 100 samples (where each sample is a vector of transcript expression estimates), the amortized cost of sampling becomes less for polee than kallisto. Approximate likelihood functions can be evaluated directly, so there is not necessarily a need to sample, but sampling is useful in some applications. For example, polee can be used as a drop-in replacement for kallisto when using sleuth [Pimentel et al., 2017] to call differential expression. Speed will be similar or faster, and this sampling technique is not subject to the limitations of bootstrap sampling, which can yield imprecise results for transcripts with a very small number of reads.

**Figure 6:**
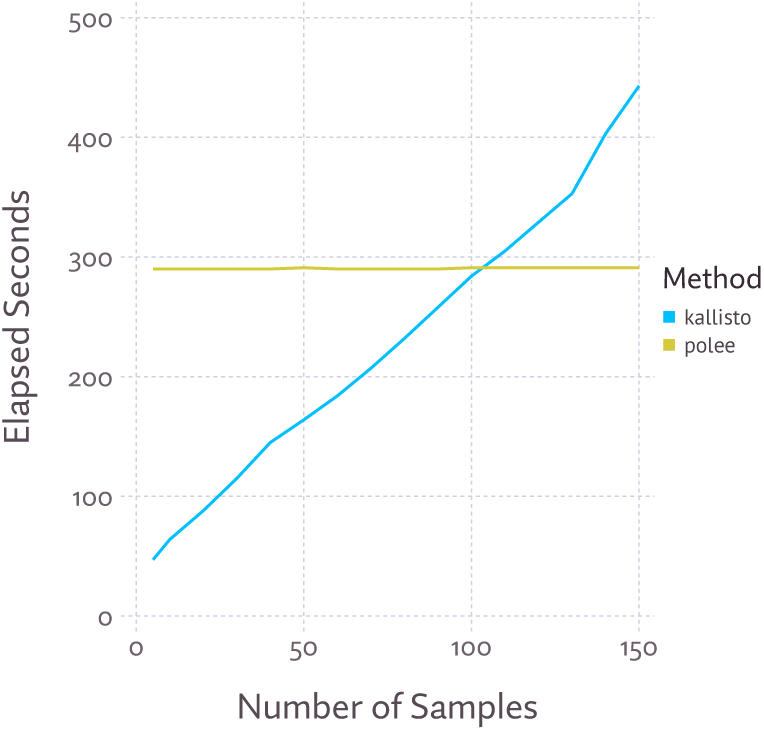
Overall run time to generate samples using kallisto (bootstrap) or polee (approximate the likelihood and sample directly from the approximation). Polee requires additional run-time up front to approximate the likelihood function, but once computed, generating samples is exceedingly efficient, with the time only increasing by about 0.69 seconds between 1 sample and 150.

### 3.3 Approximate likelihood models outperform other models in identifying differentially expressed transcripts

We expanded an analysis performed in Pimentel et al. [2017], which demonstrated superior performance when using sleuth to call differential expression, especially at the transcript level, using simulated data. We include the likelihood approximation method described here in two ways. First we developed a Bayesian regression model (see Appendix C) in TensorFlow [Abadi et al., 2016] that makes use of approximated likelihood functions directly, which is labeled “polee” in the results. Second, we generated samples from the approximated likelihood to mimic the output of kallisto, and used this as input to sleuth. This approach is labeled polee/sleuth.

The three simulations, labeled “gfr”, “isoform”, and “gcd” correspond to three sets of assumptions. The gfr simulation matches simulated effect sizes to those detected by Cufflinks [Trapnell et al., 2012] in a reference dataset. The “gcd” simulation adopts the assumption that gene expression is perturbed while holding isoform mixtures fixed, and the “isoform” simulation assumes the expression values of transcripts are perturbed independently of each other. Results from these simulations are show in Figure 7.

**Figure 7:**
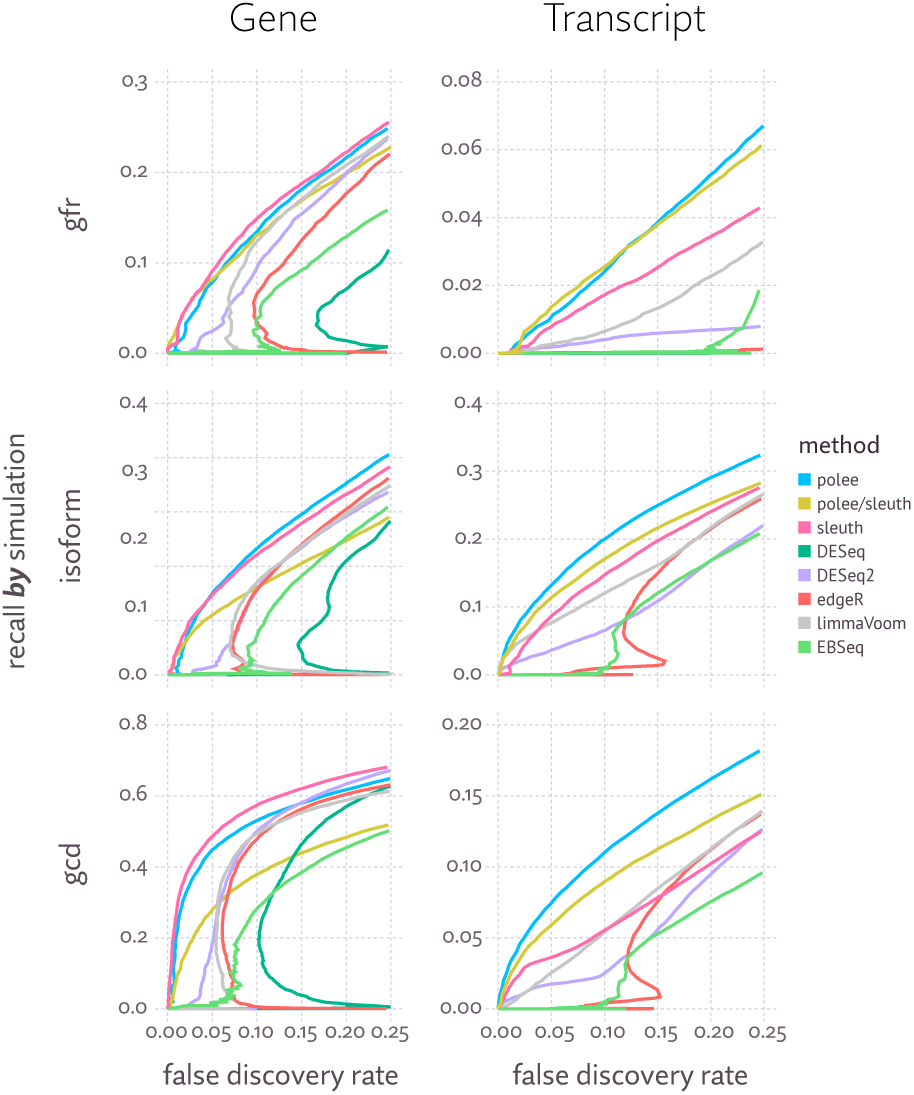
Plots of false discovery rate (i.e., the proportion of differential expression call which are incorrect) versus recall (i.e., proportion of differentially expressed features identified) when calling differential gene or isoform expression on three simulations (gfr, isoform, and gcd, explained in the text), with a variety of methods. Each line represents the aggregate FDR-recall curve across 20 replicates of the simulation, each consisting of six samples split across two conditions.

On the column to the right, we see that differential transcript expression tests show both polee and polee/sleuth with significantly improved performance over other methods (i.e., greatly increased recall at the same fdr levels). These results suggest that samples drawn in proportion to the likelihood are more informative than bootstrap samples for this task, and that most informative of all is actually including the likelihood function, or its approximation, in the model. When detecting gene-level differential expression, polee very slightly trails sleuth, which exceeds the performance of all the other methods. Oddly, using likelihood approximation samples with gene-level sleuth analysis (labeled “polee/sleuth” in Figure 7) does not yield similar performance. This may be due to sleuth’s filtering heuristics being poorly calibrated for samples from the posterior, rather that bootstrap samples.

### 3.4 Differential expression calls made with approximate likelihood are more internally consistent

Evaluating the accuracy of differential expression calls is fraught by a lack of any agreed upon ground truth. Simulations are one way around that issue, but assume the model of expression and sequencing used to generate the simulated reads is a good approximation of reality. Here we explore another option: calling differential expression with a large number of samples, then testing the ability to recover the same calls with a small number of samples. This avoids putting our faith in the verisimilitude of a simulation, but to be a reasonable proxy for accuracy it instead assumes the model converges to the correct result with enough replicates. Models can of course be both perfectly internally consistent and totally wrong, but taken together with Section 3.3 makes a case for the accuracy.

Using brain tissue data from GTEx [GTEx Consortium, 2013], we compared the same regression model using four different approaches to transcript quantification: maximum likelihood estimates, maximum likelihood with bootstrap variance estimates, posterior mean estimates generated from the likelihood approximations, and the full approximated likelihood. In addition to running regression with all 13 brain tissues, we also evaluated pairwise differential expression between a transcriptionally similar pair of tissues (hippocampus and amygdala), and a transcriptionally divergent pair (cortex and cerebellum).

Each run was compared to the same regression model run with a larger number of replicates (96 for the pairwise tests, and 1403 with all tissues). These tests were run with 10 different random subsets to draw the aggregate FDR/recall curves in Figure 8.

**Figure 8:**
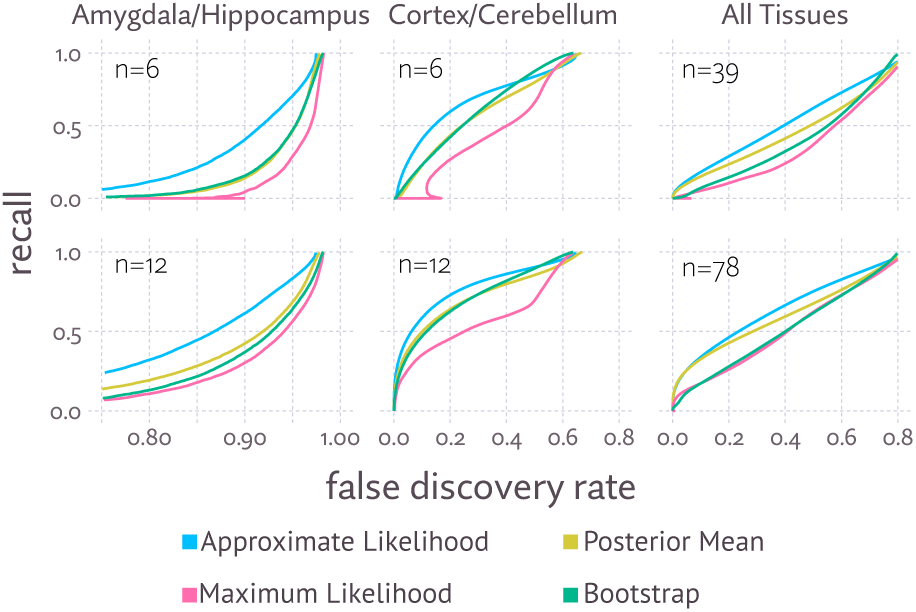
FDR/recall curves for subsets of the GTEx data, using the variants of the same regression model, with expression modeled as either maximum likelihood or posterior mean point estimates, bootstrap estimates of variance, or approximate likelihood. Note the x-axis has been adjusted for each column to show the details of the curve.

The results show broadly that the full approximate likelihood model outperforms point estimates and bootstrap estimates. Posterior mean estimates offer an improvement over maximum likelihood estimates, and appear to begin to catch up to the approximate likelihood approach when a large enough number of samples is used. An important caveat is that producing posterior mean point estimates either involves MCMC, or a variational inference, so in generating the estimates, there is little or no performance advantage to using posterior mean estimates instead of approximate likelihood. In the pairwise tests, bootstrap estimates perform similarly to using posterior mean point estimates, while underperforming when using all samples.

The amygdala versus hippocampus test shows very poor performance for all the involved methods, as the differences that do exist between these tissues are small or inconsistent, so they are only reliably detectable with a large number of samples. Nonetheless, approximate likelihood does make better use of the limited data, showing a clear improvement.

## 4 Approximate likelihood improves estimates of pairwise correlation

Large amounts of available sequencing data have led to an increasing emphasis on deciphering the functional relationships between genes. Co-expression networks are often a preliminary step towards inferring regulatory networks [Markowetz and Spang, 2007]. Constructing co-expression networks necessitates estimating pairwise correlation or covariance between genes or transcripts across samples. There are many pairs, but relatively few that are highly correlated or highly anticorrelated, so results can easily be contaminated by false positives, a particular risk with low expression genes, and pairs of similar isoforms.

Co-expression in the the GTEx data was examined by Saha et al. [2017]. To control false-positives, aggressive ad hoc filtering was done on the feature set. In addition to including only isoforms with relatively high expression (“isoforms with at least 10 samples with ≥1 TPM and ≥6 reads”), additional filters were applied for isoform variability, mappability, and many features were simply removed to maintain computational tractability. This left only 6000 genes and 9000 isoforms (for comparison, Ensembl annotates nearly 200 thousand transcripts). After filtering, a precision matrix was estimated using a graphical lasso model.

This filtering procedure reduces false-positives, but at the cost of potentially introducing false-negatives. A model affording a more principled accounting of estimation uncertainty would obviate the need for much of this ad hoc filtering.

To explore this idea, we used a more simplistic analysis of co-expression, computing pairwise Spearman correlation matrices across all annotated transcripts, with no filtering whatsoever. To see how the choice of estimate effected the results, we did this with maximum likelihood, posterior mean, bootstrap, and approximate likelihood. To avoid division by zero, and to otherwise slightly moderate the effects of zeros, we added a pseudocount of 0.1 tpm all estimates in the maximum likelihood and bootstrap samples. The uncertainty information provided by bootstrap and approximate likelihood was incorporated by computing the average Spearman correlation across 20 samples.

As with differential expression, there are no plausible gold standard estimates to compare to, so we resorted to using consistency as a proxy for accuracy. We selected one tissue from the GTEx data, cortex, consisting of 118 samples, and computed the correlation matrices. Treating this as ground truth for each respective estimate, we recomputed the matrices using random subsamples of 12 of the 118 samples, and measured the difference between each element in the matrix. This was repeated 10 times for different random subsamples. The aggregate results are plotted in Figure 9.

**Figure 9:**
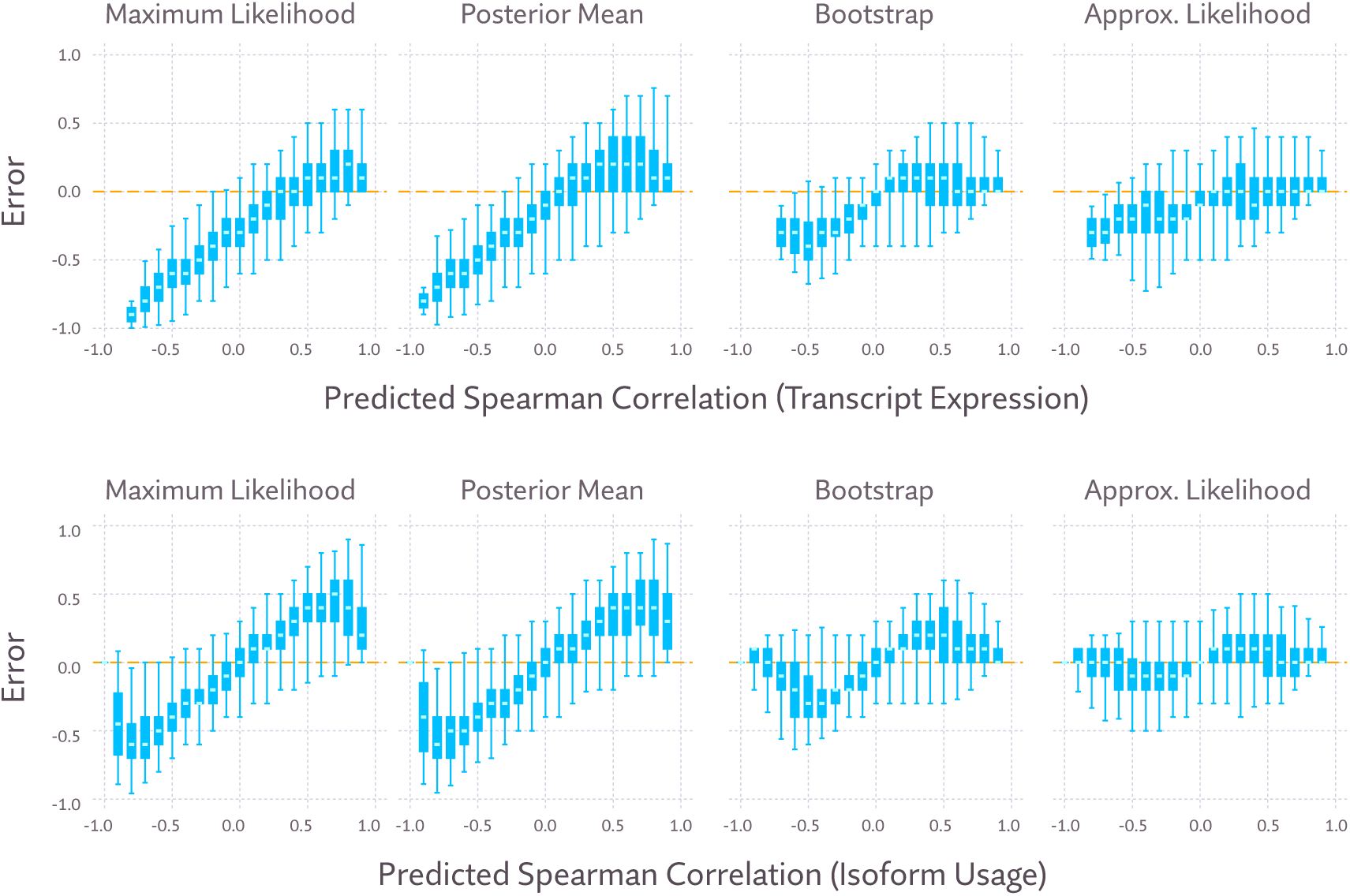
Consistency of Spearman correlation estimated using maximum likelihood and posterior mean point estimates, as well as averages across bootstrap samples, and samples from the approximate likelihood. In the upper plots, transcript expression was used, and in the lower, isoform usage (transcript expression divided by the overall expression of its gene). Error here measures the difference between estimates made using all 118 cortex samples, and estimates using a random subset of 12 samples (i.e., “true” correlation minus predicted correlation). These plots show the aggregate error across ten random subsets. Boxplots are drawn with upper and lower whiskers corresponding to the 99th and 1st percentile, respectively.

Looking first at transcript expression (Figure 9, upper plots), we see that point estimates tend to produce moderately unreliable estimates of positive correlation, and extremely unreliable estimates of negative correlation. In large part, this is remedied by using either bootstrap estimates or approximate likelihood, with the latter offering a slight improvement. When considering isoform usage (Figure 9, lower plots), point estimates are highly unreliable when measuring either positive or negative correlation. Bootstrap estimates improve this, but approximate likelihood estimates are clearly the most reliable here.

## 5 Conclusion

Here we have described how a full likelihood model of RNA-Seq transcript expression can be made tractable by approximating the likelihood function. Because including the likelihood, rather than relying on point estimates, better accounts for estimation uncertainty, differential expression calls are more reliable. The method we developed to do so, the Pólya tree transformation, is a general purpose approach to approximating sparse mixture models. Though we have confined our analysis here to showing its benefits when detecting transcript differential expression with bulk RNA-Seq, there are other possible applications. Other RNA-Seq analyses, like classification, dimensionality reduction, and coexpression could benefit from the same approach. It also presents an opportunity of performing isoform level analysis of single-cell RNA-Seq, accounting for the high estimation uncertainty where there are relatively few reads per cell.

Gelman [2016] describes the way in which statistics is sometimes used, either deliberately or otherwise, to transmute randomness into certainty as “uncertainty laundering.” The two-step process often used in RNA-Seq of first estimating, then separately modeling transcript expression can be considered a form of uncertainty laundering, but a form undertaken often out of practical necessity. We believe the method described here, a general approach to reducing the the computational demands of probabilistic RNA-Seq models, is a significant push in the direction of honest accounting of uncertainty.

## A Goodness of fit evaluated on more samples

In Section 3.1, goodness of fit of the approximation was measured by looking at distributions of p-values in null-hypothesis tests against samples from a Gibbs sampler. In that section we choose one sample to focus on arbitrarily from a mouse body map study [Li et al., 2017], but these distributions of p-values are remarkably consistent across every sample from that experiment, which we show in Figure 10.

**Figure 10:**
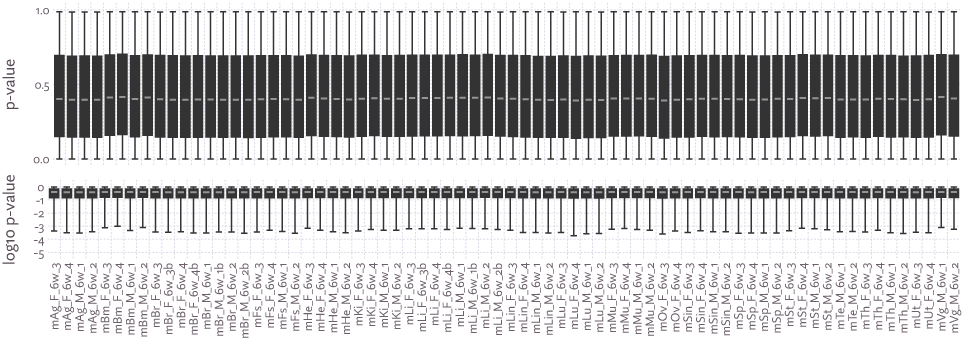
The likelihood function for every sample from [Li et al., 2017] was approximated. 1000 samples were drawn with probability proportional to the approximated likelihood, and 1000 samples were drawn using a Gibbs sampler. A Wilcoxon signed-rank test was performed for each annotated transcript, and the distribution plotted in a boxplot, drawn with upper and lower whiskers corresponding to the 99th and 1st percentile, respectively. An ideal fit would have a uniform distribution of p-values.

## B Some mathematical details of the Pólya tree transformation

In the body of the paper the Pólya tree transformation was described as a stick breaking transformation between (0, 1)^n−1^ and Δ^n−1^. Here we give a more formal definition, and generalize it somewhat by not assuming the composition sums to 1, allowing the stick being broken to be of arbitrary length.

We can then think of the transformation as between (ℝ_+_, (0, 1)^n−1^) and 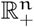, where ℝ_+_ is the set of positive real numbers. This can be thought of as mapping an initial stick length, and n − 1 break points to n stick pieces of positive length.

### B.1 Definition of the Pólya tree transformation

The Pólya tree transformation is best represented as a full binary tree, with 2n − 1 nodes, n of which are leaves. To simplify notation somewhat, assume these are assigned indices so that the root node is labeled 1, internal nodes 1, …, n − 1, and leaf nodes n, …, 2n − 1. Additionally, we assume internal nodes are numbered so that no node has a smaller index that any of its ancestors, for example, according to a pre-order traversal. Intuitively, we can think of the ith node as representing the ith break in a stick-breaking process.

The tree can then be defined by functions giving an internal node’s left and right children respectively.

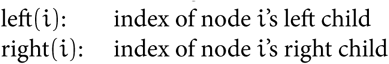

The transformation T from (u_1_, y) ∈ (ℝ_+_, (0, 1)^n−1^) to 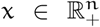 is defined in terms of *intermediate values* u_1_, …, u_2n−1_ for each node, which have the following relation,

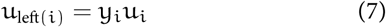

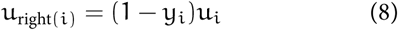

The result of the transformation is then simply the intermediate values from the leaf nodes:

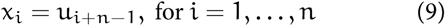

Often we are operating on the unit simplex, in which case u_1_ = 1. In these instances we will leave the u_1_ = 1 implicit and write the transformation as T : (0, 1)^n−1^ ↦Δ^n−1^.

When implemented, u_i_ values are computed by traversing the tree with any top-down traversal from the root. In the stick-breaking metaphor, intermediate values can be thought of as sizes of intermediate sticks after some number of breaks are performed. The inverse transformation can be computed by traversing the tree up from its leaves, as is done in a post-order traversal.

### B.2 Some properties of the transformation

#### **Lemma B**.**1**.

*If leaves*(i) *is the set of indexes of leaf nodes in* i*’s subtree, then*

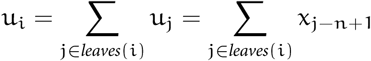

*Proof*. This simply says that each intermediate stick piece is the same length as the sum of the parts it is ultimately broken into, and is easy to see through induction.

Let i be the index of a leaf node. The subtree’s only leaf is i, so the theorem is trivially true,

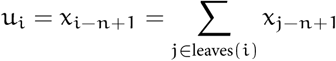

Next consider any internal node i where the theorem is true of its children. That is, where

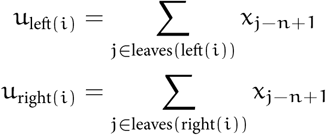

Then,

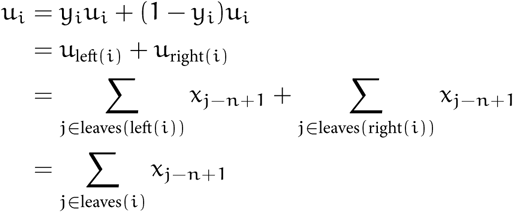

By mathematical induction, the theorem is true of all i = 1, …, 2n − 1.

□

#### **Theorem B**.**2**.

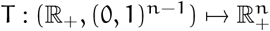 *is a bijection*.

*Proof*. For any 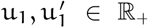, and y, y′ ∈ (0, 1)^n−1^, where 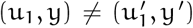, let x = T ((u_1_, y)) and 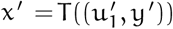.

Suppose 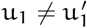, then by Lemma B.1

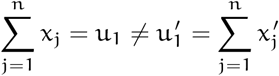

and therefore x ≠ x *′*.

Suppose 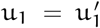, and let i be the index of the first 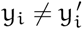. Since nodes are numbered so that ancestors have smaller indices than their children, for every ancestor j of i, 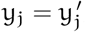, therefore 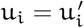. All intermediate values are positive (this follows from u_1_ being positive and elements of y residing on an open (0, 1) interval), therefore

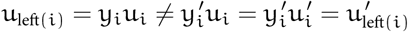

And by Lemma B.1

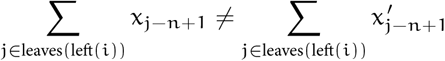

so x ≠ x*′*.

As x ≠ x *′* in either case, T is injective.

To show that T is surjective as well, consider any 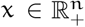.

Equations 7 and 8 can be rewritten as

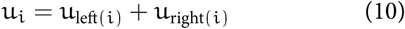

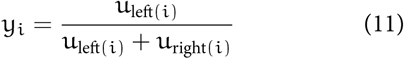

First, following Equation 9, let u_i+n−1_ = x_i_ for i = 1, …, n, thus defining leaf node intermediate values. The remaining intermediate values u_i_, for i = 1, …, n − 1 are then defined according to Equation 10, and y_i_ for i = 1, …, n according to Equation 11.

Importantly, since elements of x are positive, it follows that all intermediate values are positive, so the denominator of Equation 11 is never zero and y_i_ ∈ (0, 1).

As Equations 7, 8 are satisfied if Equations 10, 11 are, and u_1_, y are defined by the latter, it follows that T ((u_1_, y)) = x and T is surjective, and thus bijective.

□

#### **Corollary B**.**3**.

*If* u_1_ = 1 *is fixed, then the resulting transformation is a bijection between* (0, 1)^n−1^ *and* 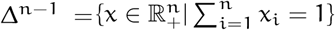.

*Proof*. A restriction of a bijection remains a bijection so we need only show that the image is Δ^n−1^. It follows directly from Lemma B.1 that the codomain of the restriction is Δ^n−1^. Furthermore, any x ∈ Δ^n−1^ has a (u_1_, y) where T ((u_1_, y)) = x, and again by Lemma B.1, u_1_ = 1, so (u_1_, y) is in the restricted domain, and thus the image is Δ^n−1^.

□

For the purposes of approximating the RNA-Seq likelihood function, we only care about this special case mapping onto the simplex.

For the approximation to be useful for computing probability densities of transformed variables, the determinant of the Jacobian must also be efficiently computable, which we show here.

#### **Theorem B**.**4**.

*Let* (u_1_, y) ∈ (Rℝ_+_, (0, 1)^n−1^), *and* J_T_ *be the Jacobian matrix of* T ((u_1_, y)). *Further, let* u_i_ *for* i = 2, …, 2n − 1 *be the intermediate values as defined in Equations 7 and 8. Then*

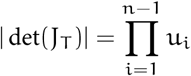

*Or in words, the absolute value of the Jacobian determinant is simply the product of the internal node intermediate values*.

*Proof*. One way to derive the determinant for J_T_ is to separate the transformation T into the composition of a number of simpler transformations. The stick breaking metaphor suggests as natural decomposition: each break can be treated as its own transformation. If T is thought of as a sequence of n − 1 breaks, it can be represented as

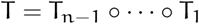

where T_i_ is the ith break and has the associated break proportion y_i_. This would let us write

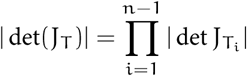

where J_Ti_ is the Jacobian matrix for T_i_.

The single break transformation

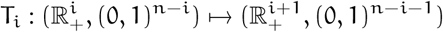

can be thought of as taking i sticks lengths, and n − i remaining breaking proportions, and breaking one of the sticks according to the first remaining proportion. Equivalently, T_i_ reflects taking one step in the tree traversal, computing u_left(i)_, u_right(i)_ from u_i_, y_i_, and then discarding these latter two values.

Incidentally, because only two values are replaced, it is not hard to see that each of these transformations is also a bijection between different n dimensional spaces.

To give a concrete example of T being decomposed into single break transformations, consider a balanced tree with n = 4 leaves, and the input u_1_ = 100, y = (0.2, 0.6, 0.9). In the following table, each row gives the input to T_i_, and the subsequent row its output. The final result is a vector in 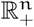.

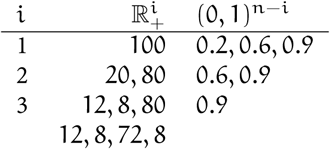

Because T_i_ only replaces u_i_, y_i_ with u_left(i)_, u_right(i)_, the Jacobian matrix is the n by n identity matrix except for a 2 by 2 submatrix so that,

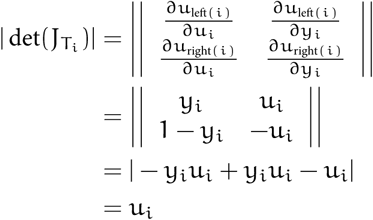

And therefore

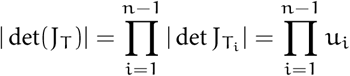

□

A Dirichlet process is a special case of a Pólya tree distribution [Ferguson, 1974]. Here we show a variation of this fact, that a Dirichlet distribution can be represented using any Pólya tree transformation, regardless of topology, if appropriately applied to Beta distributed random variables.

This is important to RNA-Seq because a multinomial distribution is often assumed when there is thought to be no read ambiguity. A multinomial distribution renormalized to form a distribution over probability vectors is a Dirichlet distribution. So this theorem says that if Pólya tree transformed Beta random variables are fit to the multinomial likelihood, nothing is lost except for a constant of proportionality. If the underlying distributions can reasonably mimic Beta distributions, then we can expect the fit to the multinomial to be very good.

**Definition B**.**5** (Hierarchical Beta distribution family). *For* n > 1, *let* T : (0, 1)^n−1^ ↦ Δ^n−1^ *be a Pólya tree transformation and* 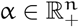.

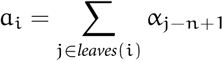

*where leaves corresponds to* T*‘s tree topology. Put in words, if we associate* α_j_ *with the* j*th leaf node* (*which is node* j + n − 1 *in our numbering scheme*), a_i_ *is the sum of* α *entries corresponding to the leaf node descendents of internal node* i.

*A random variable* X *is HierarchicalBeta*(T, α) *distributed iff* X ∼ T(Y), *where* Y = (Y_1_, …, Y_n−1_) *are independent random variables with*

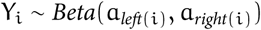

Hierarchical Beta is simply the family of distributions induced by applying a Pólya tree transformation to Beta distributed stick breaking proportions with a specifically chosen parameterization scheme. Recall that a Pólya tree distribution is a (potentially infinite) hierarchical stick breaking process [Lavine, 1992, 1994, Mauldin et al., 1992] with Beta distributed breaks. Hierarchical Beta distributions are thus a specific subset of Pólya tree distributions.

**Definition B**.**6**. *A* terminal stick break *transformation*

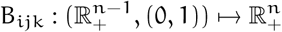

*for* 1 ≤ i ≤ n − 1 *and* 1 ≤ j < k ≤ n *is defined by*

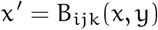

*where* x′ *is formed from* x *by removing* x_i_ *and inserting* yx_i_ *and* (1 − y)x_i_ *such that they are* 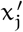 *and* 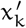, *respectively*.

*This is a bijection, and the inverse* 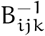, *we call an* initial stick mend. *It removes* 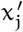 *and* 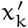 *and inserts* 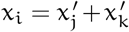, *and additionally yields* 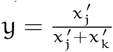.

In the proof of Theorem B.4 we described how a Pólya tree transformation can be thought of as a series of single stick break transformations, which we used to derive the Jacobian determinant. In most trees there is a choice in the order of single stick breaks, equivalent to choosing one of multiple possible pre-order traversals of the tree, each of which results in a different representation of the same Pólya tree transformation. Regardless of the ordering, there is always some final break, and this break is a terminal stick break B_ijk_ for some i, j, k.

Some notation will be useful to decompose and build up Pólya tree distributions.

**Definition B**.**7**. *If*

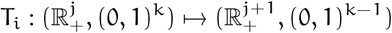

*is a single stick break transformation, as described in the proof for Theorem B*.*4, which produces one more stick length by using the first breaking p roportion, and i f* k > 1, *we can drop the last unused breaking proportion to form a new transformation we denote*

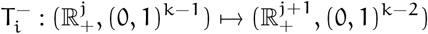

*Similarly, we can add another unused breaking proportion if* k ≥ 0, *which we’ll write as*

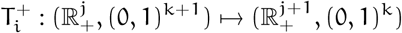

This notation lets us peel off the final break of a Pólya tree transformation. As we saw, the last break is a terminal stick break B_ijk_ for some i, j, k. So any Pólya tree transfomation T with n > 2 can be written as B_ijk_ ○ S, for a transformation

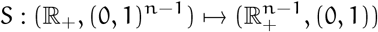

If we dispense with S’s unused breaking proportion, we have

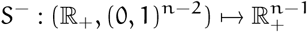

which is itself a Pólya tree transformation. Similarly, we can add another breaking proportion with the T^+^ notation to extend a Pólya tree transformation.

#### **Lemma B**.**8** (Hierarchical Beta breaking).

*Let* X ∼ *HierarchicalBeta*(T, α), *where* T *is a Pólya tree transformation. Let* 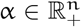, *and* Z ∼ *Beta*(a, b) *where* a + b = α_i_ *for some* 1 ≤ i ≤ n, *and let* B_ijk_ *be a terminal stick break, then*

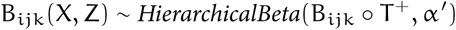

*where* α′ *is formed by removing* α_i_ *from* α *and inserting* a *and* b *such that* 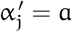 *and* 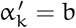.

*Proof*. The original Hierarchical Beta distribution is defined by the Pólya tree transformation T applied to n − 1 Beta distributed random variables Y_1_, …, Y_n−1_, parameterized with α as they are in Definition B.5.

Applying the terminal stick break B_ijk_ simply does one more break according to one more Beta distributed random variable Z. So the resulting distribution is defined by the the Pólya tree transformation B_ijk_ ○ T^+^ applied to Beta random variables Y′ = (Y_1_, …, Y_n−1_, Z). It only remains to be seen that Y′ follows the Hierarchical Beta parameterizations with α′.

Z is defined to trivially obey these rules, and restricting a + b = α_i_ and inserting a, b into α to form α′ preserves the parameterization of the rest of the distribution, so we can say B_ijk_(X, Z) ∼ HierarchicalBeta(B_ijk_ ○ T^+^, α′).

□

#### **Lemma B**.**9** (Dirichlet mending).

*If* X ∼ *Dirichlet*(α) *and* 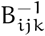 *is an initial stick mend, then*

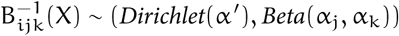

*where* α′ *is formed from* α *by removing* α_j_ *and* α_k_ *and inserting their sum so that* 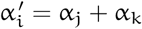.

*Proof*. This follows from two properties of the Dirichlet distribution: subcompositional invariance and amalgamation invariance (see, for example, Pawlowsky-Glahn et al. [2015] p. 121). If x ∼ Dirichlet(α), the former property says, in the narrow case needed here, that for any 1 ≤ k_1_ < k_2_ ≤ n,

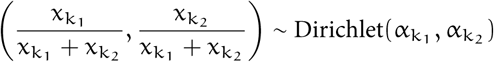

or, equivalently,

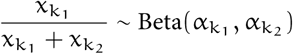

The latter property says that if {1, …, n} is partitioned into k > 1 nonempty sets I_1_, …, I_k_, then

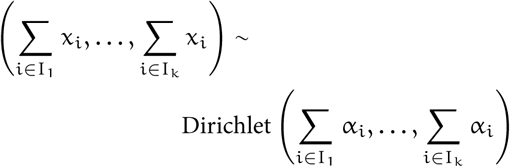

For any X ∼ Dirichlet(α), and initial stick mend B^−1^ 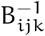 let 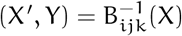.

From the definition of an initial stick mend, 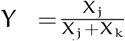. The subcompositional invariance property then gives us

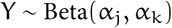

Since α′ is formed by by summing two elements of α, by the amalgamation invariance property,

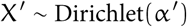

□

#### **Theorem B**.**10**.

*For any Pólya tree transformation* T : (0, 1)^n−1^ ↦Δ^n−1^ *where* n ≥ 2 *and any* 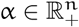

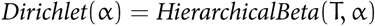

In other words, the Dirichlet distribution family and the Hierarchical Beta distribution family are exactly the same, regardless of the tree topology used for the Hierarchical Beta.

*Proof*. The proof will proceed by induction over n.

(*Base case*) Let n = 2. For any 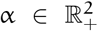 let Y ∼ Beta(α_1_, α_2_). Note that, because Dirichlet generalizes Beta, (Y, (1 − Y)) ∼ Dirichlet(α).

There is only one Pólya tree transformation T : (0, 1) ↦Δ^1^, which takes the form T(Y) = (Y, 1 − Y) and by Definition B.6, T(Y) = (Y, 1 − Y) ∼ HierarchicalBeta(T, α), so we have

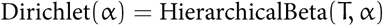

(*Inductive step*) Now assume the proposition is true for some n ≥ 2.

For any 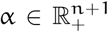 and any Pólya tree transformation T : (0, 1)^n^ ↦ Δ^n^, we can decompose T as T = B_ijk_ ○ S for transformations S and B_ijk_ where B_ijk_ is a terminal stick break for some indexes i, j, k.

By Lemma B.9, if X ∼ Dirichlet(α) then

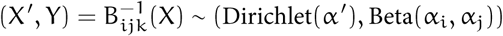

where α′ is defined as it is in Lemma B.9.

By the inductive hypothesis, X′ is also Hierarchical Beta distributed for any Pólya tree transformation onto Δ^n−1^, and since S^−^ is a Pólya tree transformation onto Δ^n−1^,

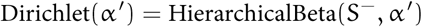

Because B_ijk_ and 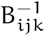 are inverses,

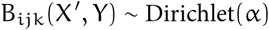

and by Lemma B.8,

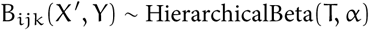

and so we have

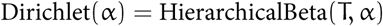

And by induction, this equality holds for all such α, T and any n ≥ 2.

□

This result can be strengthened to say that not only can any Dirichlet distribution be constructed using any Pólya tree transformation of the same dimensionality, but the Hierarchical Beta construction is the only way to do so. First, we need a simple result about random variables under bijections.

#### **Lemma B**.**11**.

*For any random variables* Y = (Y_1_, …, Y_n_) *and* 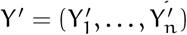, *and bijection* T, *if* T(Y) *and* T(Y′) *are identically distributed*, Y *and* Y′ *must also be identically distributed*.

*Proof*. To show the contrapositive, suppose T(Y) and T(Y′) are not identically distributed. There is then some y where P(T(Y) = y) ≠ P(T(Y *′*) = y). Since

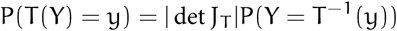

and

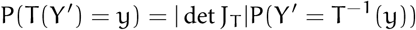

where J_T_ is the Jacobian matrix for T, it follows that

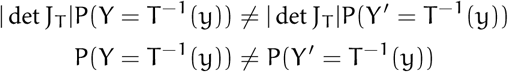

Therefore Y and Y′ are not identically distributed either.

□

Next, we can strengthen Theorem B.10.

#### **Corollary B**.**12**.

*For any Pólya tree transformation* 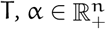, *and random variables* Y = (Y_1_, …, Y_n−1_). *If*

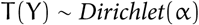

*then*

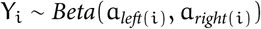

*for* i {1, …, n−1} *where* a *is defined as it is in Definition B*.*5*.

*Proof*. For any Pólya tree transformation T and α ∈ ℝ^n^, we know from Theorem B.10, that if Y = (Y_1_, …, Y_n_) are Beta distributed with the Hierarchical Beta parameterization, then

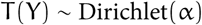

From Lemma B.11, any other random variables 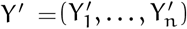 where T(Y′) ∼ Dirichlet(α) must be identically distributed to Y, and so must also be Beta distributed with the same parameterization.

□

To summarize these results: any Dirichlet distribution can be constructed using any Pólya tree transformation of the same dimensionality, and furthermore this construction is unique.

### B.3 Minimizing KL-divergence

Minimizing the KL-divergence between the approximation q and the normalized likelihood function 𝒫(r|·) (defined in Equation 3), with a set of observed reads r, uses a standard variational inference procedure, which we will briefly recapitulate here.

Recall that the approximation q can be represented as a parameterized transformation S applied to a non-parameterized distribution (i.e., the reparameterization trick). In our case, the non-parameterized distribution is simply a standard n − 1 dimensional Normal distribution N(0, I). The transformation S includes the sinh-asinh transformation, shifting and scaling, the logistic transformation, and the Pólya tree transformation T, which we collapse into a single function here for simplicity.

Explicitly, the combined transformation, for y ∈ ℝ^n−1^ is

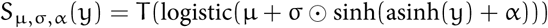

where µ, σ, α is the parameterization, ⊙ is the element-wise product, and the logistic, sinh, and asinh functions are also applied element-wise. The logistic function has the standard definition logistic(x) = (1 + exp(−x))^−1^, and T is a Pólya tree transformation. Note that T is not parameterized, since it is chosen prior to optimization, as discussed in Section 2.2.3. To simplify notation, we will combine parameter vectors as ϕ = (µ, σ, α) and use S_ϕ_ to denote the parameterized transformation.

The approximate normalized likelihood q_ϕ_ is defined as the transformation S_ϕ_ applied to a multivariate standard-normal z ∼ Normal(0, I). If p is the density function for this Normal distribution, the approximation’s density function is then

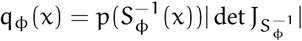

where 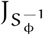 is the Jacobian matrix for the inverse S_ϕ_ transformation.

With this set up in mind, the objective function can be rewritten in a form amenable to stochastic gradient descent.

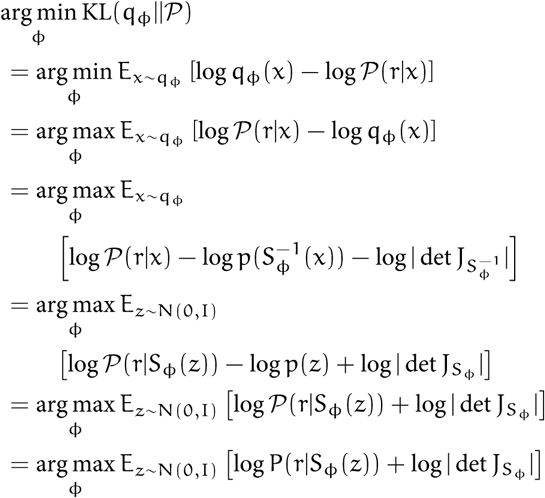

The penultimate step follows because E_z_[− log p(z)] is a constant (specifically, the entropy of a multivariate standard Normal distribution), and in the final step we replace the normalized likelihood 𝒫(·) with the actual likelihood P(r|·), since the constant of proportionality becomes an irrelevant additive constant here.

Intuitively, the objective is choosing an approximation that maximizes the expected log-likelihood, with a Jacobian term accounting for the transformation. What we are left with is an expectation than can be easily optimized through stochastic gradient descent. Each iteration of the optimization algorithm draws from N(0, I), transforms the vector, evaluates the RNA-Seq log-likelihood function, and adds to that the log of the Jacobian determinant of the transformation, computing the gradients with respect to ϕ. In practice we use Adam [Kingma and Ba, 2014] to update ϕ estimates each iteration, and 500 iterations, with each iteration estimating gradients using 6 Monte Carlo samples of z.

## C Regression Model

Differential expression is determined throughout the paper using a probabilistic linear regression model on log-expression, but with the additional layer of approximate likelihood.

In defining the model we will use the following notation:

m number of samples in the experiment

n number of transcripts

k number of factors/columns in the design matrix

X m by n matrix of log-expression values

A m by k design matrix

W k by n matrix of regression coefficients

µ length n log-expression bias vector

σ length n transcript standard deviation vector

r some representation of the RNA-Seq reads

**1** ones vector (length m in all uses below)

The basic regression model is defined by,

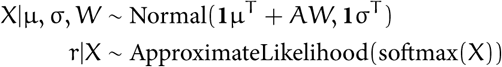

The softmax functions transforms the log-expression to relative expression. There are some subtleties of this move noted below. The bias vector µ has a Normal prior with a large variance.

Apart from the inclusion of the extra layer r|X, treating the reads as observed and expression values as unobserved, this is a fairly standard log-linear model of gene expression, similar to the one described by Smyth [2004], for example.

### Prior on regression coefficients

Operating under the prior expectation that most transcripts are not differentially expressed, W is given a sparsity inducing prior. Elements of W_ij_ have the prior given by,

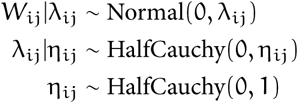

This is a version of what Bhadra et al. [2017] describe as the “Horseshoe+” prior.

### Modeling variance

Transcript expression standard deviation σ is shrunk towards standard deviation estimates of transcripts with similar expression.

To do this we choose h = 15 “knots” at expression values, equally spaced between the minimum and maximum, and compute weights u_jℓ_ between every transcript j and knot ℓ, using the squared difference between the knot’s expression value and the transcript expression averaged across samples. These weights are pre-computed using initial approximate posterior mean expression estimates, obtained by applying the approximation’s transformation to a zero vector. Each knot has associated parameters α_l_, β_l_.

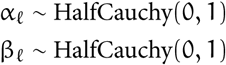

Per-transcript standard deviation is Inverse-Gamma distributed with parameters computed as weighted sums of these.

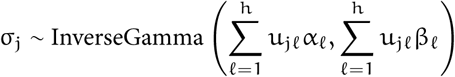

### Scale penalization

Because RNA-Seq measures relative expression, detecting changes is absolute expression involves additional assumptions. Various normalization schemes have been proposed to overcome this [Bullard et al., 2010]. Other methods avoid the problem by looking for compositional changes [McGee et al., 2019]. Similar to the normalization schemes, our model attempts to find changes in absolute expression by assuming most transcripts or genes are maintaining relatively constant expression (or constant lack of expression).

Because of the the softmax transformation, log-expression values are not on a fixed scale, which is to say for a given sample 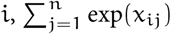 is not constrained to sum to any particular number. How relative scales between samples are inferred determines how changes in transcript composition are interpreted as changes in expression.

Because of the sparsity inducing prior, higher probability is achieved when fewer transcripts are differentially expressed. If this assumption is reasonable relative scaling will be arrived at automatically so as to minimize differences between samples. Of course, there are circumstances when this assumption fails, particularly in the presence of large changes in the overall quantity of RNA between groups of cells.

In practice we found it helpful for inference to additionally penalize these exponent sums from straying too far. To find reasonable values, we use initial posterior mean estimates to estimate relative scales that minimize differential expression of highly expressed transcripts, then include an additional Normal prior in the model on these exponent sums, centered on the scale estimates. This is simply one approach to putting an informative prior on relative scale.

### Inference

The model is fit using variational inference implemented in TensorFlow. Computing the Pólya tree transformation efficiently, particularly many different transformations with different tree topologies, using tensor operations is not straightforward. To avoid this issue, we implemented a custom TensorFlow operation in C++ which computes transformations in parallel on CPUs. Currently, there is no GPU version of this operation, so while running inference on GPUs greatly speeds it up (for example, running a regression on 96 samples took 44 minutes with a GPU, and 168 minutes without), computing the Pólya tree transformation remains the bottleneck, and an obvious target for further improvements in efficiency.

When calling differential expression with this model, effect size is always considered explicitly. Because the model is continuous, the posterior probability of no change in expression is zero. We instead look for transcripts or genes with a sufficiently large posterior probability of the effect size being above some threshold. Unlike most null-hypothesis test models which typically (though, not necessarily) adopt a null hypothesis of zero change, there is an extra threshold that must be chosen: the minimum effect size. This is a slight complication, but has the advantage of being up front about what is considered a potentially interesting effect.

